# The buoyancy of cryptococcal cells and its implications for transport and persistence of *Cryptococcus* in aqueous environments

**DOI:** 10.1101/2024.05.20.595024

**Authors:** Isabel A. Jimenez, Piotr R. Stempinski, Quigly Dragotakes, Seth D. Greengo, Lia Sanchez Ramirez, Arturo Casadevall

**Affiliations:** Department of Molecular and Comparative Pathobiology, Johns Hopkins University School of Medicine, Baltimore, Maryland, USA; Department of Molecular Microbiology and Immunology, Johns Hopkins University Bloomberg School of Public Health, Baltimore, Maryland, USA

**Keywords:** environmental pathogens, halocline, pathogen transmission, marine mammals, public health, water, wildlife health

## Abstract

*Cryptococcus* is a genus of saprophytic fungi with global distribution. Two species complexes, *C. neoformans* and *C. gattii*, pose health risks to humans and animals. Cryptococcal infections result from inhalation of aerosolized spores and/or desiccated yeasts from terrestrial reservoirs such as soil, trees, and avian guano. More recently, *C. gattii* has been implicated in infections in marine mammals, suggesting that inhalation of liquid droplets or aerosols from the air-water interface is also an important, yet understudied, mode of respiratory exposure. Water transport has also been suggested to play a role in the spread of *C. gattii* from tropical to temperate environments. However, the dynamics of fungal survival, persistence, and transport via water have not been fully studied. The size of the cryptococcal capsule was previously shown to reduce cell density and increase buoyancy. Here, we demonstrate that cell buoyancy is also impacted by the salinity of the media in which cells are suspended, with formation of a halocline interface significantly slowing the rate of settling of cryptococcal cells through water, resulting in persistence of *C. neoformans* within 1 cm of the air-water interface for over 60 min and *C. gattii* for 4-6 h. Our data also showed that during culture in yeast peptone dextrose media (YPD), polysaccharide accumulating in the supernatant formed a raft that augmented buoyancy and further slowed settling of cryptococcal cells. These findings illustrate new mechanisms by which cryptococcal cells may persist in aquatic environments, with important implications for aqueous transport and pathogen exposure.

**Importance:** Cryptococcosis is a major fungal disease leading to morbidity and mortality worldwide. *C. neoformans* is a major fungal species of public health concern, causing opportunistic systemic infections in immunocompromised patients. *C. gattii* was traditionally a tropical pathogen, but in the 1990s emerged in the temperate climates of British Columbia and the Pacific Northwest United States. Outbreaks in these areas also led to the first host record of cryptococcosis in free-ranging cetaceans. *C. gattii* is particularly concerning as an emerging fungal pathogen due to its capacity to cause clinical disease in immunocompetent patients, its recent spread to a new ecological niche, and its higher resistance to antifungal therapies. Our research defines characteristics that influence transport of cryptococci through water and its persistence at the air-water interface, which improve our understanding of mechanisms for cryptococcal aqueous transport and persistence.

## Introduction

*Cryptococcus* is a genus of environmental fungi with global distribution. Two members, *C. neoformans* and *C. gattii*, belong to species complexes that cause pulmonary and neurologic infections in humans, domestic animals, and wildlife.^1–7^ *C. neoformans* is considered ubiquitous in the environment, while *C. gattii* is endemic in tropical and subtropical regions. However, since the 1990s, *C. gattii* has emerged as the cause of outbreaks in humans and animals in the temperate regions of British Columbia, Canada and the Pacific Northwest of the United States, raising new questions regarding the ecological niches, persistence, and spread of this fungal pathogen.^7, 8^ The historical record and epidemiological factors surrounding active *C. gattii* outbreaks suggest that water plays a key role in cryptococcal dispersal and propagation. Clinical isolates have been proposed to trace their origin to Northern Brazil,^9^ having been anthropogenically transported by shipping routes^10^ and later carried onto land by tsunami-related floods into coastal forests.^11^ This capacity for aqueous transport means that cryptococcal species may have the potential for global spread via ocean currents.

Cryptococcal species previously identified from water samples include *C. gattii*^12–14^ and *C. neoformans*^15^ demonstrating that water may be a reservoir of pathogenic cryptococci, as well as other species such as *C. albidus*, *C. laurentii*, and *C. humicolus.*^16–18^ Cryptococcal cells have been identified in freshwater, brackish water, and seawater, from coasts to deep sea trenches, at water surfaces, and from biofilms in municipal water systems.^12–25, 26^ *C. gattii* outbreaks in the Pacific Northwest have also resulted in the first cryptococcosis cases in free-ranging marine mammals.^1, 8, 27^ Cryptococcal infections in humans and land animals involve inhalation of dry aerosols in the form of spores and/or desiccated yeasts from terrestrial environmental reservoirs, such as soil, wood, dust, and dried avian guano.^28–30^ However, these recent marine mammal infections demonstrate that inhalation of cryptococci suspended in water could also be a viable mode of natural infection, and thus liquid droplets and aerosols present an understudied mode of respiratory exposure for susceptible individuals. Defining factors that influence the survival and persistence of cryptococci in aquatic environments is therefore pertinent to understanding disease transmission.

A major virulence factor of *Cryptococcus* is the capsule, comprised of branched polysaccharides anchored at the cell wall and radiating outwards with decreasing density.^31, 32^ Prior work from our laboratory demonstrates that larger capsules decrease cell density and thus increase buoyancy, potentially serving as a flotation device and facilitating dispersion through water.^33^ In the current study, we further analyzed the contribution of the capsule to buoyancy. In addition, because cryptococci in soils have small capsules,^34^ we hypothesized that the capsule may not be the primary mechanism by which cryptococcal cells remain buoyant when washed from land to sea, and sought to evaluate additional mechanisms by which cryptococci could persist in water, with a particular focus on persistence at the air-water interface. Here we report that cryptococci utilizes a variety of mechanisms to remain suspended in water and that aquatic environments can support buoyancy of cryptococcal cells.

## Materials and Methods

### Yeast strains, culture conditions, and media

Frozen stocks of *C. neoformans* (H99 (ATCC 208821) and acapsular *cap59* deletion mutant (C536 derived from B-3501 parental strain^35, 36^) and *C. gattii* (environmental isolate WM161 (ATCC MYA-4562), and feline clinical isolate NIH 409^37^) were inoculated into liquid Yeast Peptone Dextrose (YPD) media (BD Difco, Sparks, MD) and incubated in a culture rotator (37 rpm) at 30 °C for 48 h. Confluent cultures were streaked onto solid YPD media (BD Difco), incubated at 30 °C for 48 h, and stored at 4 °C until inoculation into liquid culture. Unless otherwise indicated, all four strains were utilized in each experiment, and cells were cultured for 1-2 days at 30 °C in liquid YPD media. Minimal media (MM) was prepared as previously described.^38^ Pacific Ocean seawater (SW) (Imagitarium, Petco, San Diego, CA) and live Nutri-Seawater**^®^** Aquarium Saltwater (LSW) (Nature’s Ocean, Fort Lauderdale, FL) were purchased from commercial vendors. Where indicated, seawater was filter-sterilized using a 0.22 µM filter (Sigma Aldrich, Burlington, MA).

### Cell imaging and measurements

Cells were photographed with India ink counterstaining on an Olympus AX70 microscope (Olympus America, Melville, NY) at 20 X or 40 X magnification, with a QImaging Retiga 1300 camera using QCapture software (QImaging, Burnaby, British Columbia, Canada).^39^ Cell diameter and cell body diameter were measured from at least 50 cells per condition in Fiji^40^ and capsule volume was calculated.^39^ Capsule:body volume ratio was calculated by dividing capsule volume by cell body volume.

### Capsule formation during seawater incubation

Baseline cell measurements were taken. To induce capsular enlargement, 50 µL of culture was inoculated into 5 mL of MM and incubated for 3 d. Remaining YPD culture samples were separated into two aliquots, centrifuged at 2300 g for 4 min, resuspended in PBS or filter-sterilized SW, and incubated for 3 d. Cell measurements were repeated.

### Percoll density gradient

Cells were washed and resuspended in PBS. A working solution of Percoll^®^ (MilliporeSigma) was prepared to a final osmolality of 1.0914 g/mL, as previously described.^33^ Colored polyethylene Density Marker Beads (DMB) (Cospheric, Santa Barbara, CA) were used as density standards (green, 1.02 g/cc; orange, 1.04 g/cc; violet, 1.06 g/cc; dark blue, 1.08 g/cc; red, 1.09 g/cc; medium blue, 1.13 g/cc). A volume of 50 µL (1 x 10^7^ cells) of each culture or 20 µL of each DMB was added to 13 x 51 mm polypropylene centrifuge tubes (Beckman Coulter, Sykesville, MD) containing 3 mL of Working Percoll Solution (WPS). Tubes were balanced with WPS and centrifuged using an Optima TLX tabletop ultracentrifuge (Beckman Coulter) with TLA 100.3 fixed angle rotor at 40,000 rpm at 25 °C for 30 min.^33^ Tubes were photographed using a Nikon D3000 DSLR camera under uniform light conditions.

### Halocline formation and sodium chloride specific gravity standard curve

Halocline interfaces form in nature whenever freshwater flows onto seawater, such as in estuaries and caves, resulting in vertical stratification of the fluids by density, with low density freshwater forming a relatively stable surface layer. To demonstrate halocline formation as a function of differences in specific gravity of the suspension media and cuvette media, phenol red indicator (Sigma Aldrich) was dissolved in PBS or SW. A volume of 200 µL of each solution was added to PMMA cuvettes (Plastibrand, Germany) containing 3 mL of either PBS, SW, or LSW and cuvettes were photographed. In a separate experiment, a standard curve of sodium chloride was prepared (**Supplemental Table 1**). An overnight YPD culture of H99 was washed once in PBS and resuspended in a solution of phenol red indicator dye (PBS-PR), and 200 µL of cell suspension was added to each cuvette. Photographs were taken within 1 min. To demonstrate dynamic persistence of the halocline layer, cells of strain WM161 were suspended in phenol-red dyed PBS, layered onto SW in a conical tube and agitated while video was captured.

### Buoyancy assays

Cells were washed once and resuspended in PBS, MM, or filtered SW. Cuvettes were prepared with 3 mL of PBS, MM, or filtered SW and 200 µL of cell suspension (1-3 x 10^7^ cells) was gently added to the top of each cuvette. Control cuvettes received 200 µL of PBS with phenol red. Settling was photographed at intervals.

### Suspension and passive settling

Cuvettes containing 1.5 mL of YPD media were lined up in a dark box and 1.5 mL of confluent cell culture was added, for a final cell concentration of 2-3 x 10^8^ cells/mL. Cells were resuspended using gentle manual pipetting and photographed at intervals. This suspension experiment was also repeated using cells heat-inactivated in a water bath at 60 °C for 1 h.

### Settling rate calculation

A cuvette was marked in millimeter intervals and photographed under identical conditions to experimental cuvettes. Adobe Photoshop was used to add digital measurement lines and the distance between the water surface and the upper border of suspended cells was measured. Cell settling was calculated as the rate of displacement from the surface over time.

### Phenol-sulfuric acid assay

An overnight culture of strain 409 was passively settled for approximately 18 h at room temperature before collection of 500 µL of the translucent upper layer. Concurrently, a 500 µL sample of confluent overnight culture was collected. Samples were diluted (1:100), vortexed to disrupt large polysaccharide aggregates, and centrifuged at 2300 g for 4 min. The supernatant was saved, and the pellet was washed twice and resuspended in water. A phenol-sulfuric acid assay to detect total polysaccharides was performed as previously described.^41^ Absorbance was measured at 490 nm, and readings normalized to background readings from control wells of water. Polysaccharide concentration (µg/mL) was calculated using the standard curve and normalized to the cell count of each sample.

### Immunocytochemistry

Samples (50 µL) were washed once, pelleted and resuspended in 200 µL of a 10 µg/mL solution of 18B7 (IgG1) murine monoclonal antibody (mAb)^42^ (Unisyn Technologies) in 1% BSA-PBS blocking buffer, to label glucuronoxylomannan (GXM) polysaccharide. Samples were incubated at 4 °C overnight with gentle agitation. Cells were washed and incubated at room temperature for 1 h with 2.5 µg/mL goat anti-mouse IgG Alexa-Fluor 488 secondary antibody (ThermoFisher), and 5 µg/mL Uvitex 2B (Polysciences Inc., Warrington, PA) to label cell wall chitin, in 1% BSA-HBSS. Simultaneously, a 5 mL overnight culture of strain 409 was passively settled and the upper layer was collected. To preserve polysaccharide architecture, no washes were performed. A 50 µL sample of material was diluted in 150 µL of HBSS and incubated overnight with 18B7 (10 µg/mL final concentration) at 4 °C with gentle horizontal agitation. The sample was then incubated at room temperature for 1 h with 5 µg/mL of goat anti-mouse IgG Alexa-Fluor 488 secondary antibody and 5 µg/mL of Uvitex 2B. Samples were imaged on a Leica THUNDER Live Cell and 3D Confocal Microscope at 63 X (oil objective). Minimum and maximum brightness of each image channel was set uniformly for all images and composites created in Fiji. In a separate experiment, material from the upper layer of a settled 409 culture was incubated at room temperature for 1 h with 5 µg/mL of Uvitex 2B and 10 µg/mL of 18B7 mAb directly conjugated to fluorophore Oregon Green 488, according to manufacturer instructions (ThermoFisher), and then imaged at 63 X (oil objective).

### Mucicarmine stain

Cytology slides of strain 409 material were air-dried, fixed in 100% ethanol for 2 min and stained with Mayer’s Mucicarmine Method for Mucin and *Cryptococcus* kit (PolyScientific R&D Corp.) according to manufacturer instructions and imaged at 100 X (oil objective). Mucicarmine binds to low-density negatively-charged acidic mucin, staining it reddish purple, while nuclei are stained black.

### Lipid droplet induction and lipid quantification

To induce lipid droplet formation, cells of strains H99 and 409 were cultured in plain YPD media or YPD supplemented with 4 mM oleic acid (Sigma Aldrich). A buoyancy assay was performed using polystyrene cuvettes (Globe Scientific). To quantify lipid, culture samples (200 µL) were incubated for 5 min with 5 µg/mL Uvitex 2B, then incubated with 25 µL of 1:1 DMSO:PBS for 1 min to permeabilize cells, followed by 125 µg/mL of Nile Red in acetone for 5 min to stain neutral lipids.^43, 44^ Samples were imaged at 63 X (oil objective). Nile Red fluorescence intensity was measured from at least 50 cells per strain and condition using Fiji. Readings were normalized to background fluorescence intensity from blank spaces.

### Specific gravity by refractometry

Specific gravity (SG) is the ratio between the density of a compound and the density of pure water at 4°C (1.000 g/cm^3^) and is proportional to the salinity of a liquid. SG of each media type was measured using a salinity refractometer.

### Statistical analysis

Statistical analyses were performed using GraphPad Prism 10.1.1. To assess for differences in the rate of settling, strains and conditions were compared by simple linear regression or nonlinear regression using a one-phase decay model, as indicated. To evaluate the significance of differences in capsule size, cell body size, and capsule:body volume ratio, a Kruskall-Wallis test with Dunn’s multiple comparisons testing was performed. To compare fluorescence intensity of samples stained with Nile Red, outliers were identified via ROUT (Q=1%) and excluded from analysis, and unpaired t-tests were used to compare groups. To compare polysaccharide concentrations between samples, a one-way ANOVA with Sidak’s correction for multiple comparisons was performed.

## Results

### Cell density and cell dimensions for four *Cryptococcal* strains

We measured cell densities of four strains (H99, *cap59*, WM161 and 409) using a Percoll gradient and found differences in cell density and density heterogeneity (**Figure 1A**). Capsule:body volume ratio was calculated (**Figure 1B**). Strain-specific differences were present in cell body size (**Figure 1C**), capsule size (**Figure 1D**), and capsule:body volume ratio (**Figure 1E**). Notably, the average cell body radius of *cap59* cells was significantly larger than for strains H99, WM161, and 409 (P<0.0001), which would increase cell density. Strain 409 had a significantly smaller average cell body radius than strains *cap59*, H99, and WM161 (P<0.0001) and larger average capsule radius than strains H99 or WM161 (P<0.0001), contributing to a larger capsule:body volume ratio than strains H99 or WM161 (P<0.0001).

**Figure 1.**
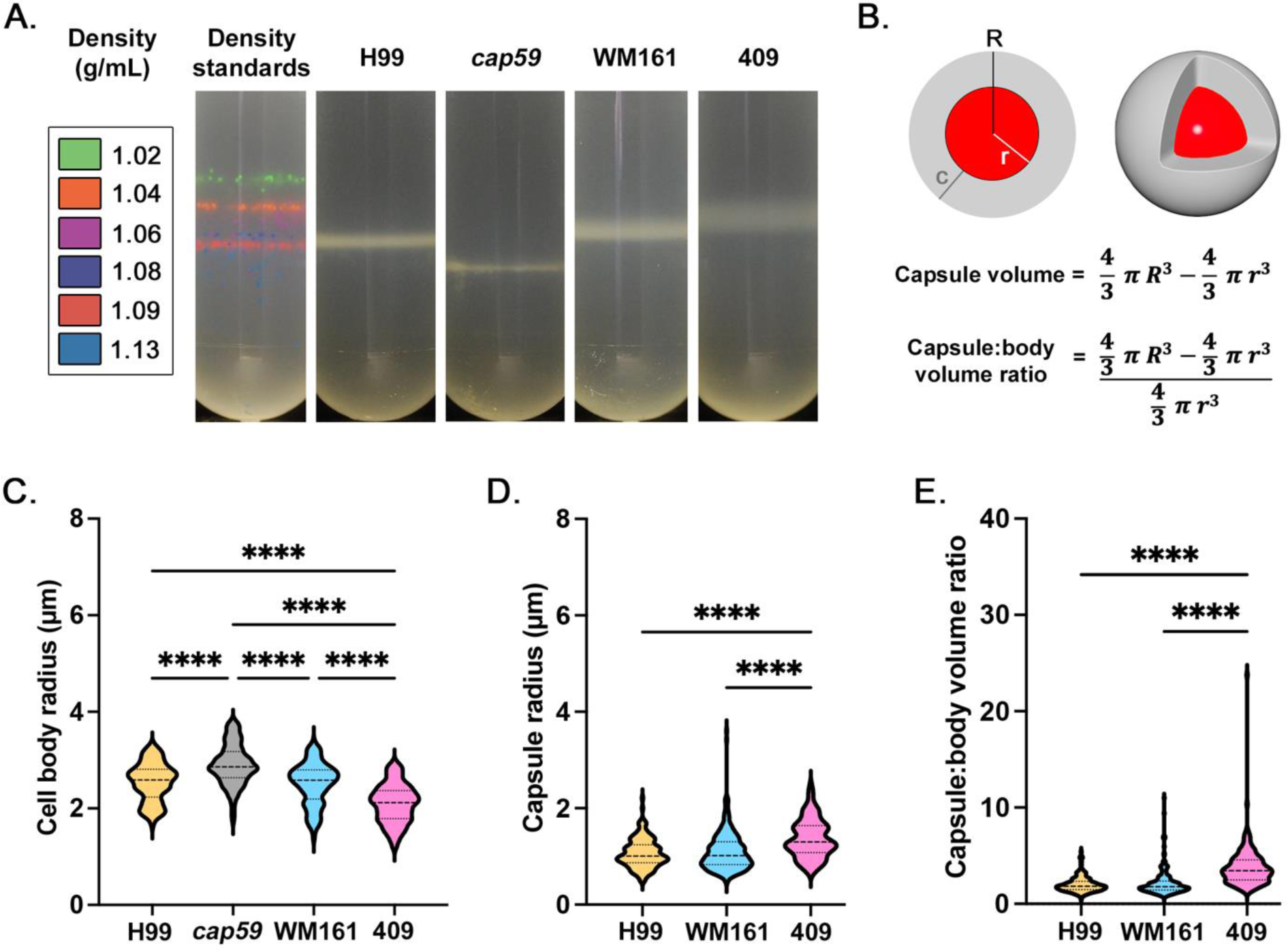
Comparison of cell density and measurements of four strains of *Cryptococcus*. *C. neoformans* strains H99 and acapsular mutant *cap59*, and *C. gattii* strains WM161 and 409, were cultured overnight in Yeast Peptone Dextrose (YPD) media. Results represent at least three independent experiments per strain and condition. **A)** Density was evaluating using a Percoll density gradient, in comparison to a standard of density marker beads. Strain *cap59* had the highest cell density (1.13 g/mL), followed by H99 (1.08-1.09 g/mL). Strains WM161 (1.06-1.085) and 409 (1.04-1.07 g/mL) had wider and less dense bands. **B)** Diagram of a cryptococcal cell, with cell body (red) and capsule (gray). Measurements were taken of total radius (R), cell radius (r), and capsule radius (c). The capsule volume was calculated by subtracting the volume of the cell body from the volume of the entire cell. The ratio between capsule volume and cell body volume was calculated by dividing the capsule volume by the cell body volume. **C)** At baseline, cell body size varied significantly between strains, with *cap59* having a larger cell body and 409 having a smaller cell body. **D)** Baseline capsule radius was significantly larger for 409 compared to H99 and WM161. **E)** Strain 409 had a significantly larger baseline capsule:body volume ratio than H99 and WM161.

### Effect of aqueous culture conditions on cell growth and capsule size

Cells were inoculated into MM, PBS, or filtered SW. As expected, MM incubation induced significant capsule enlargement in all encapsulated strains compared to overnight YPD culture (P<0.0001), while no change in capsule size was observed following SW incubation for strains H99 (P=0.1606), WM161 (P=0.6451) or 409 (P>0.9999) (**Figure 2A**). Incubation in PBS resulted in significant increase in capsule size for *C. gattii* strains WM161 (P<0.0001) and 409 (P=0.0005), but not for *C. neoformans* strain H99 (P>0.9999). Strain WM161 and strain 409 had very similar appearance; representative images of strains H99 and 409 are shown (**Figure 2B**).

**Figure 2.**
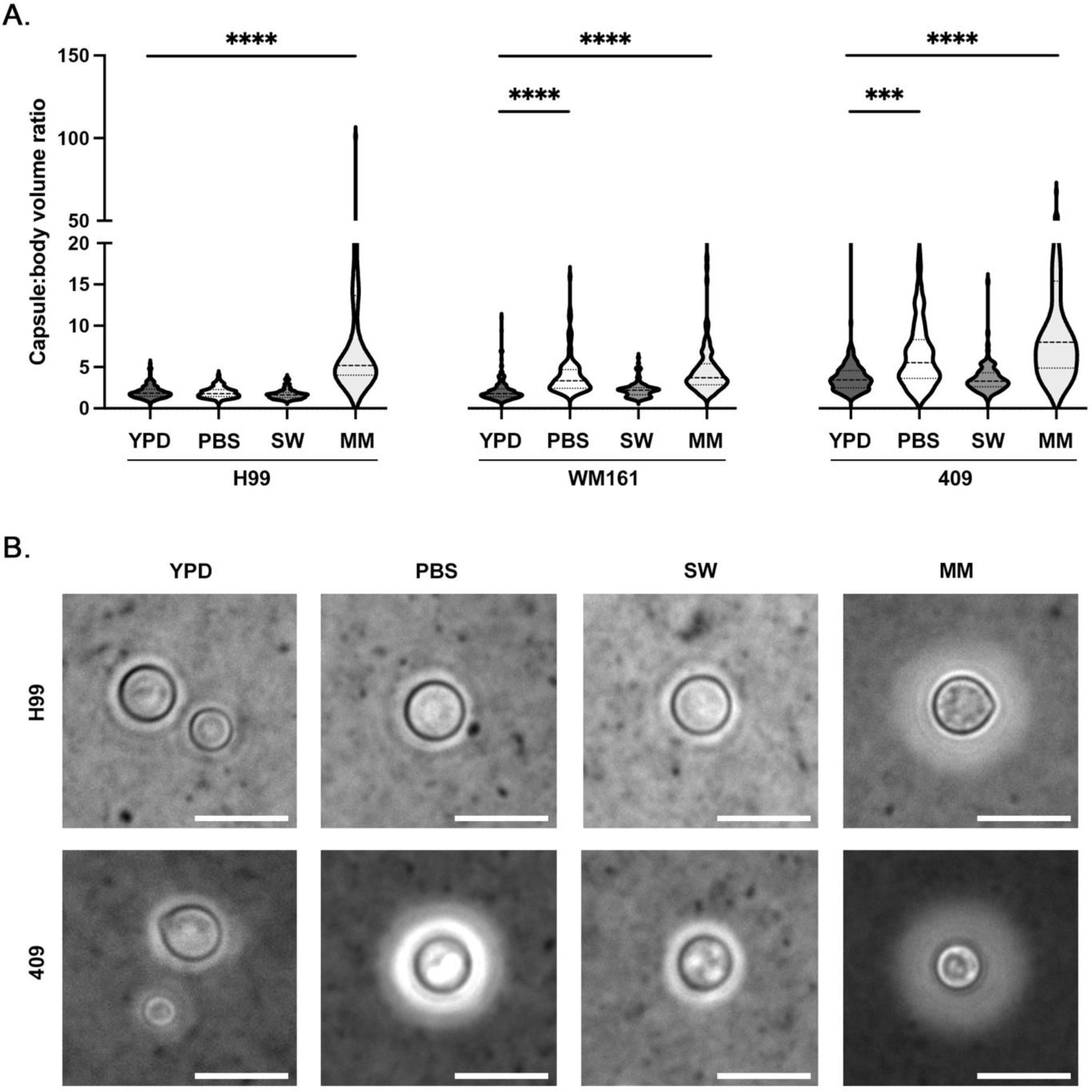
Effect of incubation in different media types on capsule growth and cell survival. *C. neoformans* strains H99 and *C. gattii* strains WM161 and 409 were cultured overnight in YPD media, washed once in phosphate buffered saline (PBS), and then incubated for 3 d in PBS, filtered seawater (SW), or minimal media (MM). Capsules and cell bodies were measured, and capsule:body volume ratio was calculated. Results represent two independent experiments per strain and condition. **A)** Incubation in SW did not induce significant capsular growth in any strain (H99, P=0.1606; WM161, P=0.6451; 409, P>0.9999). Both *C. gattii* strains had larger average capsule:body volume ratios after PBS incubation compared to baseline (WM161, P<0.0001); 409, P=0.0005), while the average capsule:body volume ratio of strain H99 did not change in response to PBS incubation (P>0.9999). As expected, incubation in MM resulted in capsular growth in all strains (P<0.0001). **B)** Representative microscopy images with India Ink counterstaining showing relative differences in capsule size between H99 and 409 strains under four media conditions. Strain H99 showed no change in capsule size in response to incubation in PBS or SW, while strain 409 developed a larger capsule following PBS incubation. Strain WM161 was similar to strain 409 in appearance. Scale bar = 10 µm.

### Specific gravity by refractometry

Specific gravity (SG) of each media type was measured by salinity refractometry (**Table 1**). SG of cell suspensions, each at a concentration of 1 x 10^8^ cells/mL in each media type, were also measured; SG of each cell suspension was unchanged from the SG of plain media.

**Table 1:**
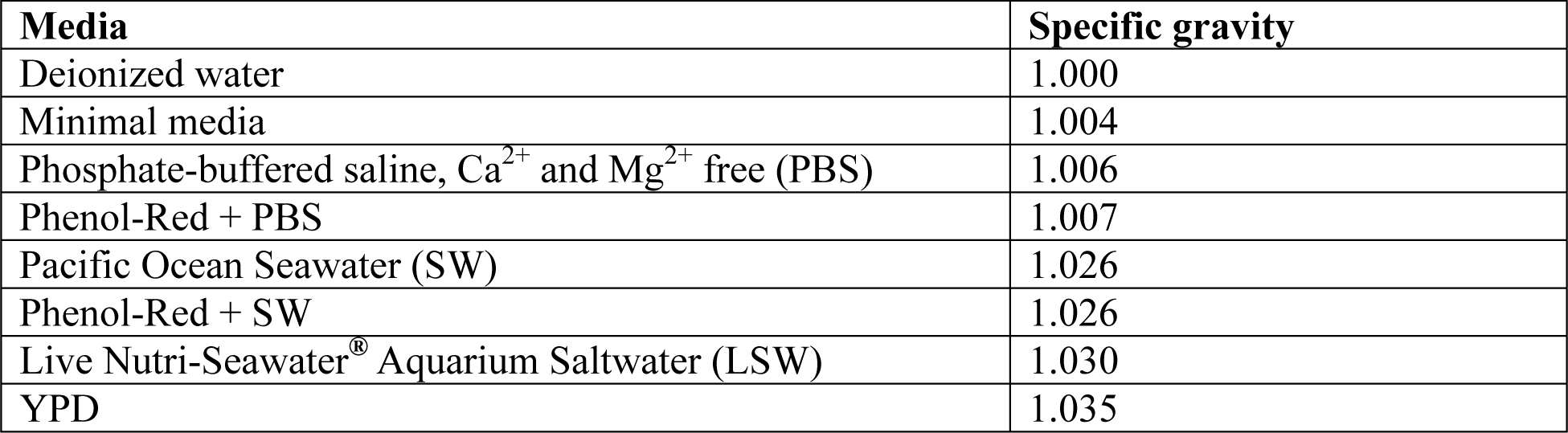
Specific gravity of media by salinity refractometry.

### Experimental halocline interface formation

We experimentally illustrated the formation of a halocline interface, with and without the presence of cryptococci. When the difference in SG (ΔSG) between the suspension media (SG_1_) and the media in the cuvette (SG_2_) is negative, a halocline forms, as illustrated by addition of PBS with phenol-red (PR) indicator dye (PBS-PR) onto SW (**Figure 3A**). Conversely, when ΔSG ≥ 0, the liquids rapidly mix. Even very small differences in SG are important, as demonstrated by halocline formation after addition of SW-PR (SG = 1.028) atop LSW (SG = 1.030) (ΔSG = −0.004), while no halocline formed after addition of PBS-PR (SG = 1.007) to a column of PBS (SG = 1.006) (ΔSG = 0.001). To evaluate the influence of ΔSG on halocline size and to illustrate the effect of the halocline on suspended cells, we prepared a sodium chloride standard curve and added cells of strain H99, suspended in PBS-PR. As ΔSG increases, the halocline interface becomes narrower, trapping cells close to the water surface (**Figure 3B**).

**Figure 3.**
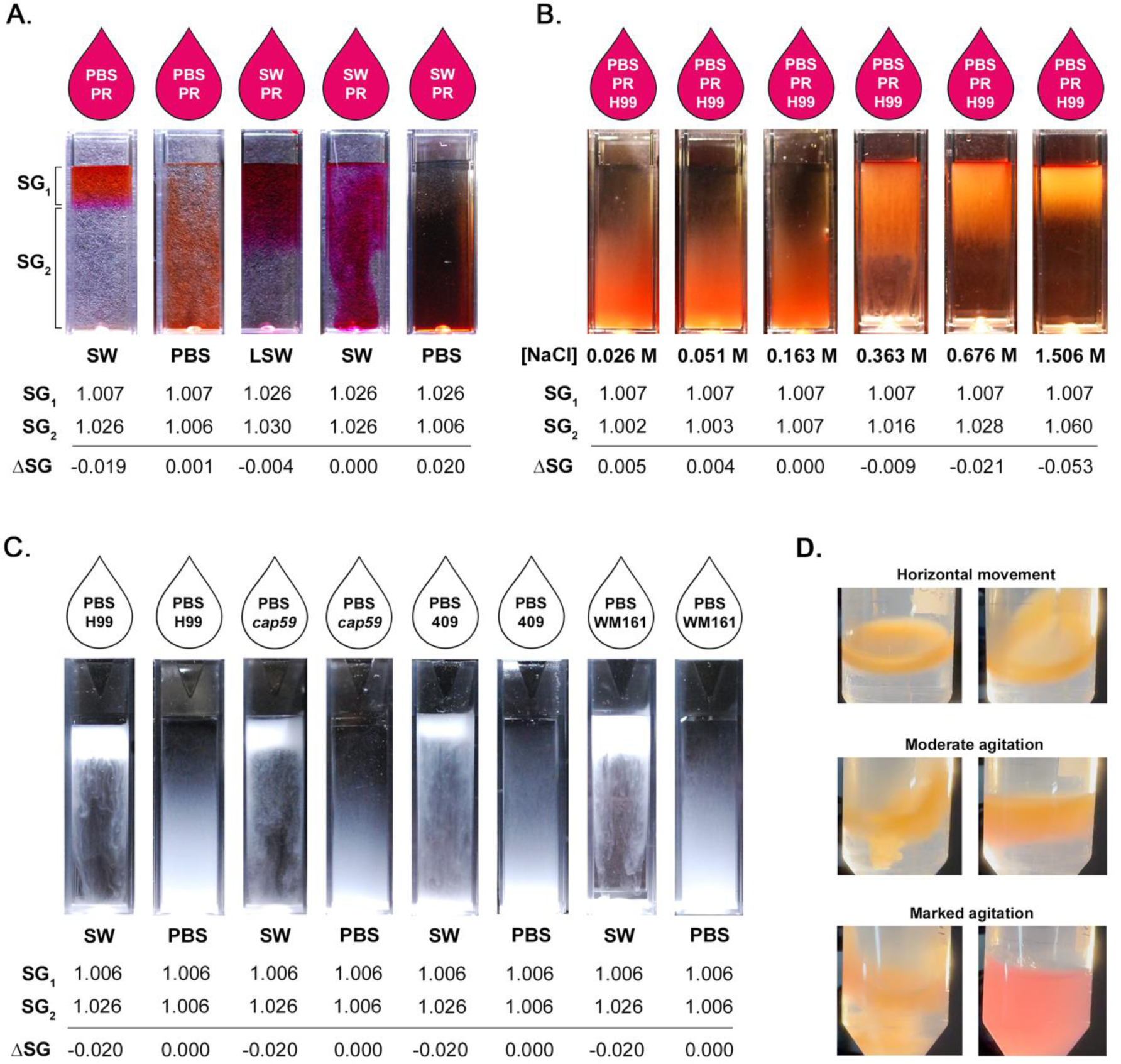
Experimental halocline formation results in suspension of cryptococci at the air-water interface. Specific gravity (SG) is the ratio between the density of a compound and that of pure water at 4°C (1.000 g/cm^3^) and is proportional to salinity. When two liquids are layered, the difference in specific gravity (ΔSG) determines if layers vertically stratify to form a halocline interface. **A)** If ΔSG < 0, a halocline forms, as illustrated by the addition of PBS with phenol-red (PR) indicator dye (PBS-PR) onto a column of Pacific Ocean seawater (SW). Even very small differences in SG result in halocline formation, as shown by addition of SW-PR (SG = 1.028) to seawater from a different source (LSW; SG = 1.030). Conversely, when ΔSG ≥ 0, no halocline forms and the liquids rapidly mix. **B)** Strain H99 was suspended in PBS-PR and added onto cuvettes containing serial concentrations of NaCl, illustrating the impact of ΔSG on the size of the halocline. **C)** Strains H99, *cap59*, WM161 and 409 were suspended in PBS, and then layered onto either PBS or SW. When cells were suspended in PBS and added to SW, a halocline interface temporarily trapped cells in the top layer. This effect is seen even in the acapsular *cap59* mutant. Conversely, when cells grown under identical conditions were suspended in PBS and added to PBS, the cells dispersed rapidly, demonstrating the marked impact of halocline layer formation on buoyancy. **C)** When cryptococcal cells of strain WM161 were suspended in PBS and added to a conical tube of SW, cells became trapped in the halocline interface, which remained stable throughout gentle tilting, horizontal movement of the tube, or gentle agitation. The stratification was disrupted only with marked agitation, as observed by the rapid color change of the indicator dye.

To demonstrate that halocline formation was sufficient to suspend cryptococci at the air-water interface regardless of capsule size, we cultured cells in YPD media, suspended cells in PBS, and added them on top of columns of PBS or filtered SW. For all four strains, cells were buoyant when added to a column of SW, whereas the same cells sank rapidly when layered onto PBS (**Figure 3C**); this pattern was conserved despite the absence of a capsule in the *cap59* mutant and varying capsule sizes in the other strains. We further illustrated that cells remained suspended above the halocline under dynamic conditions, only mixing once the tube was vigorously shaken (**Figure 3D**).

### The presence of a halocline interface slows the rate of cryptococcal settling

A buoyancy assay time-course was performed from 5 min to 4 h. Cells suspended in PBS were layered onto cuvettes of filtered SW, resulting in halocline formation (**Figure 4A; Supplemental Figure 1**). Concurrently, cells suspended in PBS or filtered SW were added to the top of cuvettes containing PBS; under both conditions, cells were carried by the suspension media to the bottom of the cuvette. In the presence of a halocline, cells of strains H99, WM161 and 409 became trapped at the upper 1 cm of the cuvette for over 60 min. Under the two combinations that incorporated filtered SW, cells of the acapsular mutant strain *cap59* exhibited marked macroscopic clumping and adherence to the walls of the PMMA cuvette; this granulated appearance persisted for hours. Rates of cell settling were calculated for each strain and condition except *cap59*, for which only settling in PBS could be assessed. Cells of strains H99 (**Figure 4E**), 409 (**Supplemental Figure 2**), and WM161 (**Supplemental Figure 2**) settled significantly slower in the presence of a halocline (P<0.0001). In the absence of a halocline, cells settled fastest when suspended in PBS and added to PBS (SG 1.006), while cells settled at an intermediate rate when suspended in filtered SW and added to PBS (final SG 1.009) (**Figure 4E**; **Supplemental Figure 2**). Further, strain-specific differences in rate of settling were apparent, each strain having a unique exponential decay function (P<0.0001) (**Figure 4F**).

**Figure 4.**
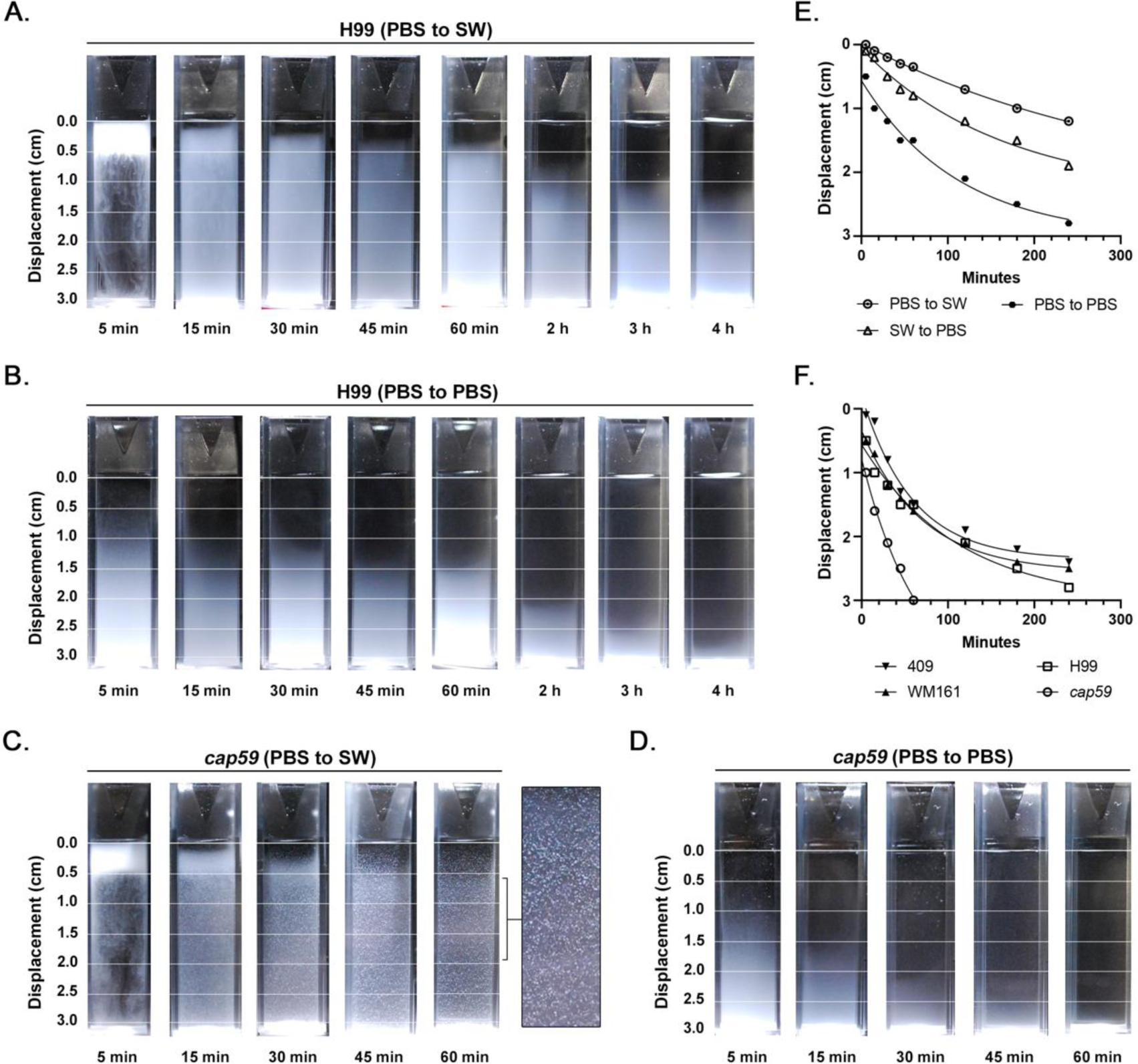
The halocline delays settling of cryptococci in seawater. Cryptococci of strains H99, *cap59*, WM161 and 409 were suspended in PBS or filtered seawater (SW) and added to the top of cuvettes containing either PBS or SW. The rate of cell settling was assessed over 4 h by measuring displacement (cm) from the top of the cuvette. Images and graphs shown are representative of two independent experiments. **A)** Cells of strain H99 suspended in PBS and added to SW were initially suspended at the halocline interface and then moved out of the halocline over time. **B)** Cells of strain H99 suspended in PBS and added to PBS settled more rapidly. For images of all strains and conditions, see **Supplemental Figure 1. C)** Acapsular *cap59* cells suspended in PBS and added to SW exhibited marked clumping and adherence to the side of the PMMA cuvette. **D)** Acapsular *cap59* cells suspended in PBS and added to PBS did not exhibit clumping. **E)** Media type significantly impacted the rate of cell settling (P<0.0001). Cell settling was slower in the presence of a halocline interface. In the absence of a halocline, cell settling was proportional to the final specific gravity (SG) in the cuvette, with cells suspended in SWF and added to PBS (SG = 1.009) exhibited an intermediate rate of settling, while cells suspended in PBS and added to PBS sank most rapidly (SG = 1.006). This figure panel shows data for strain H99, with trends representative of all strains; for strains WM161 and 409, see **Supplemental Figure 2. F)** Strain-specific differences in rate of settling were observed for all media conditions. This panel shows the rate of settling of cells suspended in PBS and added to PBS. For strain-specific rates of settling for the other two media conditions, see **Supplemental Figure 2**.

### Strain-specific rates of passive settling

To assess dynamics of cryptococcal settling through water in the absence of a halocline, cell cultures were resuspended in cuvettes and allowed to passively settle. The acapsular mutant *cap59* settled completely by 60 min. Strains H99, WM161, and 409 were each incompletely settled by 6 h, with both *C. gattii* strains settling slower than H99 (**Figure 5A**). Upon observation after 26.5 h, all strains had settled, but WM161 and 409 still demonstrated two distinct layers, with a translucent upper layer and an opaque lower layer; this finding was much more prominent for strain 409. Although transient, we also visualized a translucent upper layer after 1.5-2 h of settling in strain H99, although without clear delineation between layers. This translucent upper layer was notably absent in *cap59*. Rates of passive settling, as determined by the slope from addition of cells until settling (1 min to 6 h) using simple linear regression, were significantly different between strains (P<0.0001, F = 50.30) (**Figure 5B**). In a separate trial, cells were cultured overnight in YPD and cell concentration was adjusted using fresh YPD media to match that of the prior experiment before the suspension assay was repeated. The overall relationship between rate of strain settling (*cap59* > H99 > WM161 > 409) was unchanged. For the *cap59* mutant, the rate of settling was the same between the two media conditions. However, all encapsulated strains settled significantly faster in the refreshed media compared to the original media (**Supplemental Figure 3**). No differences in rate of settling were observed between live and heat-inactivated cells.

**Figure 5.**
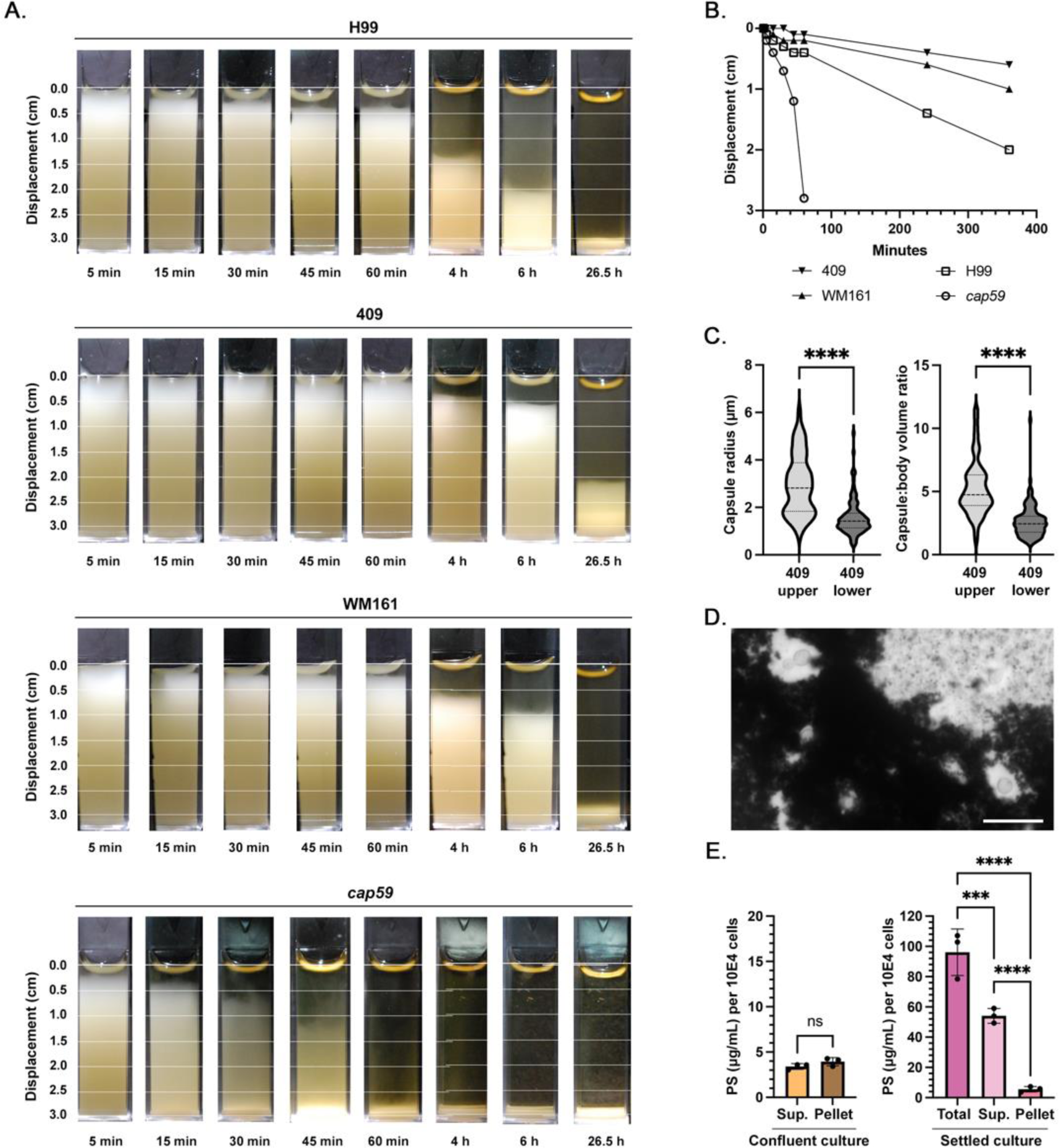
Passive settling times for *C. neoformans* and *C. gattii* are strain-specific. *C. neoformans* strains H99 and *cap59*, and *C. gattii* strains WM161 and 409, were grown overnight in liquid YPD culture, resuspended in cuvettes, and allowed to passively settle while photographs were taken at intervals between 5 min and 6 h, and then again at 26.5 h. Results of **A-C** represent two independent experiments. **A)** The *cap59* acapsular mutant settled most rapidly, with all cells settled after 60 min. Strains WM161 and 409 settled slower than H99, and even when fully settled, exhibited two distinct layers. **B)** Based on linear regression, the rate of passive settling was significantly different between strains (P<0.0001). **C)** Cells from the upper layer of a settled strain 409 culture had significantly larger capsule radii (P<0.0001) and capsule:body volume ratio (P<0.0001) compared to cells from the lower layer. **D)** Attempts to perform microscopy of the upper layer of a settled strain 409 culture using standard protocols for India ink counterstaining revealed marked clumping of material with cells with large capsules entrapped inside. Scale bar = 20 µm. **E)** Using a phenol-sulfuric acid assay, the polysaccharide concentration of a confluent strain 409 culture was quantified and compared to that of the upper layer of a settled strain 409 culture, demonstrating that passive settling significantly enriched polysaccharide content in this upper layer.

### Polysaccharide in culture affects cell settling by forming rafts entrapping cells

Samples of strain 409 cultures were allowed to passively settle overnight and the translucent upper layer was collected by gentle manual pipetting. Cells in the upper layer were less concentrated (1-3 x 10^7^ cells/mL) than in the lower layer (1-2 x 10^9^ cells/mL) across two independent replicates. Cells in the upper layer had significantly larger capsule radii (P<0.0001) and capsule:body volume ratios (P<0.0001) than cells in the lower layer (**Figure 5C**). The upper layer was largely comprised of acellular material, which induced clumping of India Ink (**Figure 5D**) and formed structures entrapping cells. On phenol-sulfuric acid assay, polysaccharide concentrations in strain 409 were significantly higher in samples from the upper layer of a settled 409 culture compared to a confluent culture (P<0.0001) (**Figure 5E**). In addition, the combined polysaccharide concentration of the supernatant and cell fractions decreased by 38% after washing (P=0.0003) demonstrating that a significant amount of polysaccharide is lost during the wash steps.

We performed immunocytochemistry to visualize the capsule. When washes were performed between incubation steps, cells of strains WM161 and 409 had more irregular and diffuse capsule margins compared to strain H99, with strain 409 also having dimmer fluorescence (**Figure 6A**). When immunocytochemistry was performed without wash steps on material collected from the upper layer of a settled 409 culture, we observed large aggregates with branched structures and strong 18B7 fluorescence, suggesting that these are heavily composed of polysaccharides of the same type as the capsule (**Figure 6B, 6C**). Furthermore, a subset of cells in proximity to these aggregates had absent, dim, or irregular capsular binding of 18B7. When samples of this material were stained with mucicarmine, we visualized a large proportion of cells with threads of capsular material extending to adjacent cryptococci, and lacy patches of pale purple-staining material between cells (**Figure 6D**).

**Figure 6.**
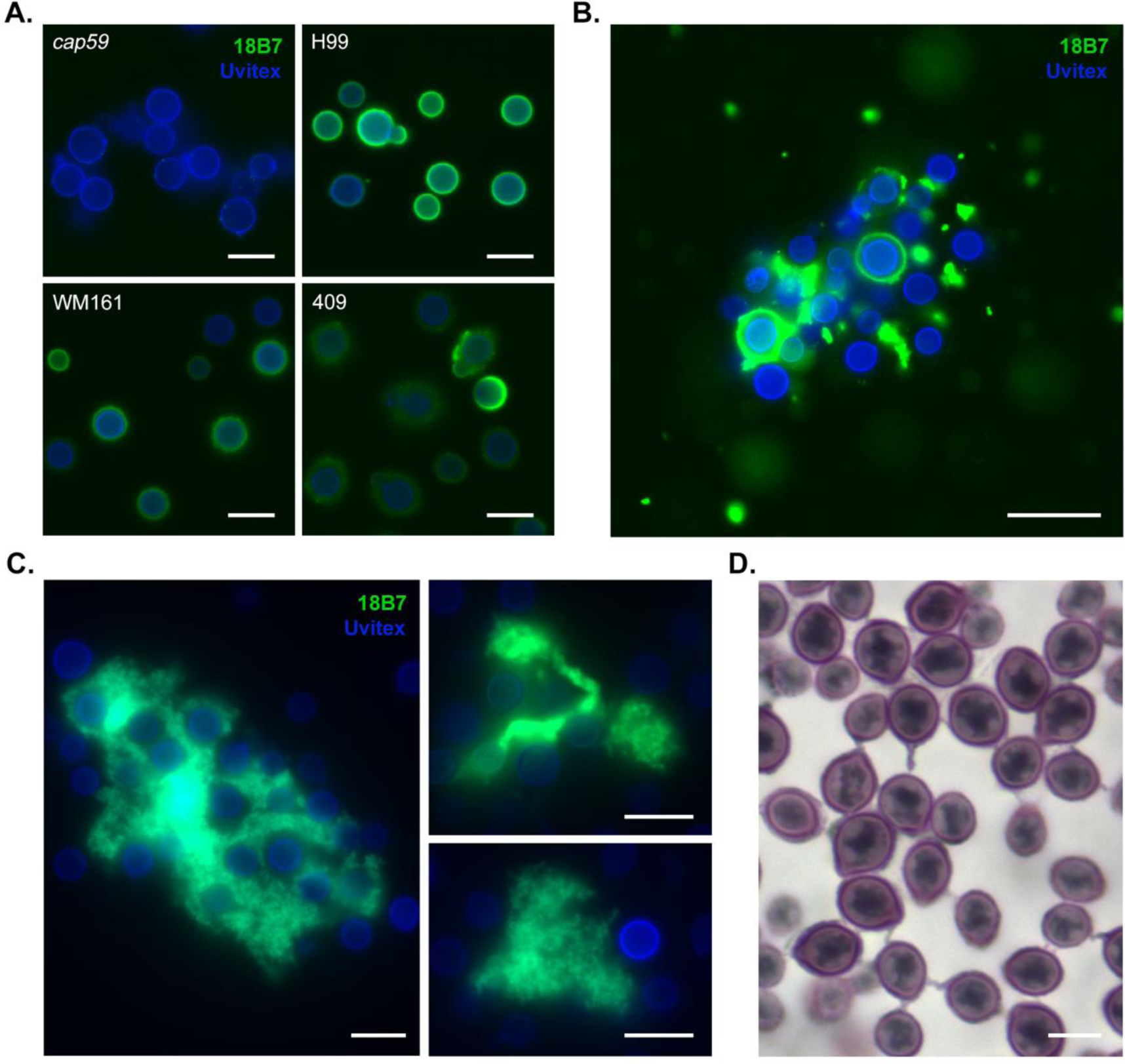
Polysaccharide in the cell supernatant forms large aggregates and entraps cells of *C. gattii*. **A)** Immunocytochemistry was performed on YPD cultures of strains H99, *cap59*, WM161 and 409 using 18B7 murine monoclonal anti-capsular antibody (green) and Uvitex 2B for cell wall chitin (blue). Cells were washed between primary and secondary antibody incubation steps per standard immunocytochemistry protocols. We observed that cells of strains WM161 and 409 had more irregular and diffuse capsule margins compared to strain H99, with strain 409 also having dimmer fluorescence. The acapsular *cap59* mutant is unable to export polysaccharide to the capsule and exhibits no fluorescence. Scale = 10 µm. **B)** When a strain 409 culture was allowed to settle and immunocytochemistry was performed on this material, without wash steps, we observed large aggregates of material with strong 18B7 fluorescence, suggesting that they are heavily composed of polysaccharides of the same type as the capsule. Furthermore, a subset of cells in proximity to these aggregates had absent, dim, or irregular capsular binding of 18B7, suggesting that an abundance of free extracellular polysaccharide can sequester antibody. Scale = 20 µm. **C)** Intricate branched structures were observed within large clumps of polysaccharide entrapping cells. Scale bar = 10 µm. **D)** A large proportion of cells showed threads of capsular material extending to adjacent cryptococci, entrapped in lacy patches of pale purple-staining material, consistent with polysaccharide. Mucin is stained reddish purple, nuclei are stained black. Scale bar = 5 µm.

### Strain-specific differences in lipid content and assessment of lipid contribution to buoyancy

After culture in plain media, strain 409 had significantly higher mean (P<0.0001) and maximum (P<0.0001) fluorescence intensity than strain H99 (**Figure 7C, 7D**). For strain H99, mean (P<0.0001) and maximum (P<0.0001) fluorescence intensity were higher after incubation in oleic acid-supplemented media compared to plain media (**Figure 7E**). Strain 409 grown in oleic acid-supplemented media had significantly higher maximum fluorescence intensity (P=0.0002) but no significant difference in mean fluorescence intensity (P=0.1230) compared to cells grown in plain media (**Figure 7F)**. There was no significant difference in the rate of cell settling for 409 cells between growth conditions. Although H99 cells cultured in oleic-acid supplemented media appeared to settle slightly slower than cells in plain media at certain timepoints, differences in overall rate of cell settling between culture conditions did not reach statistical significance at the α = 0.05 level (P=0.0700) (**Figure 7A, 7B**).

**Figure 7.**
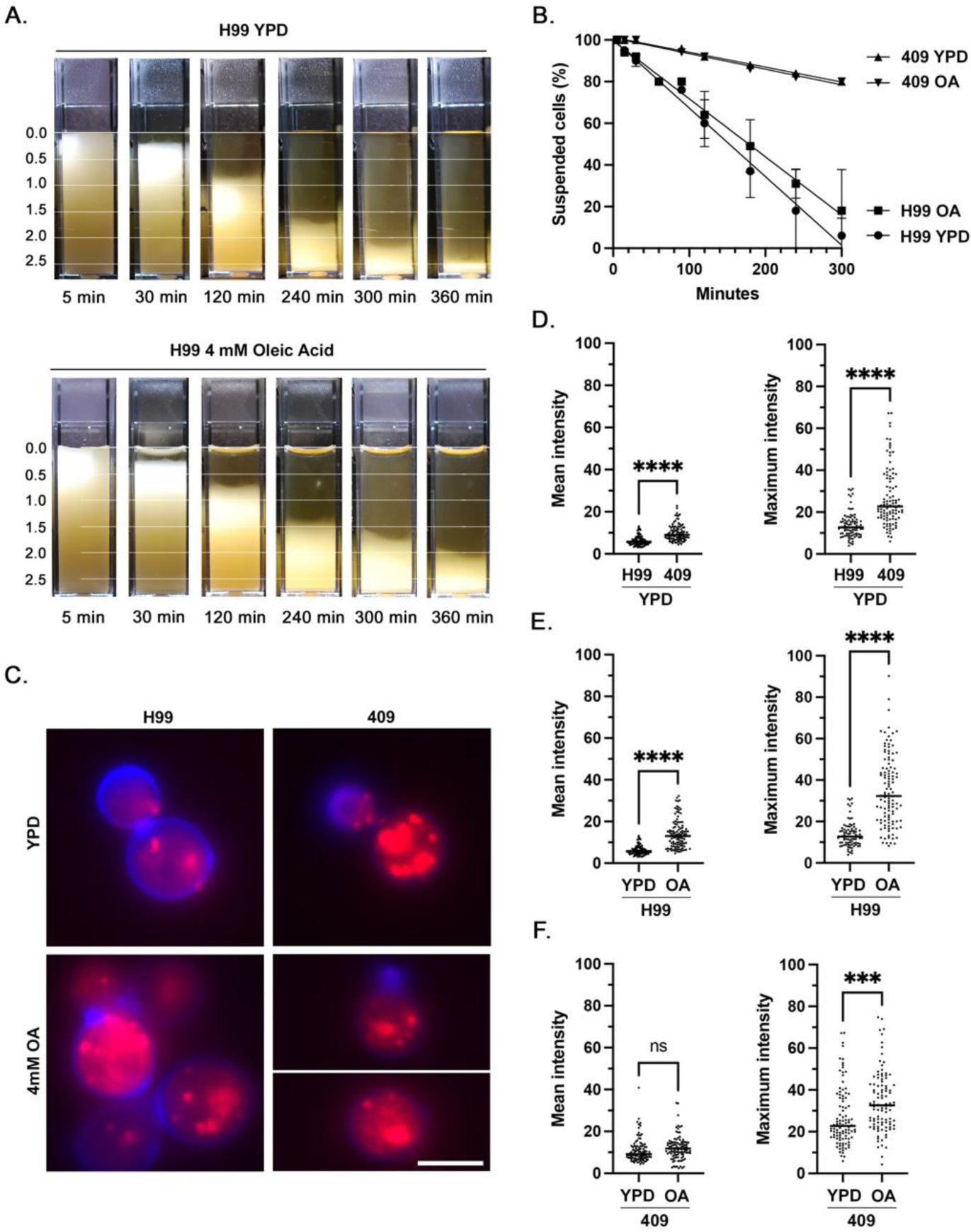
Effect of oleic acid supplementation on intracellular lipid accumulation and cell buoyancy in *C. neoformans* strain H99 and *C. gattii* 409. **A)** Cuvettes of H99 cells were grown overnight in plain YPD media or YPD media supplemented with 4 mM Oleic Acid, resuspended and allowed to passively settle. Photographs were taken at 10 timepoints between 5 min and 360 min; shown are a subset of timepoints. Data represents the results of two independent experiments. **B)** Rate of cell settling by simple linear regression. Strain 409 cultures settled significantly faster than strain H99 cultures (P<0.0001), while culture conditions did not result in a significant difference in the rate of settling for strain 409 (P=0.4539) or strain H99 (P=0.0700). **C)** Immunofluorescence images representative of relative fluorescence of neutral lipid (Nile Red, red) and cell wall chitin (Uvitex, blue) in strains H99 and 409, with and without oleic acid supplementation. Scale bar = 5 µm. **D)** After overnight culture in plain YPD media, strain 409 cells had a higher mean and maximum fluorescence intensity of neutral lipid compared to strain H99. **E)** After overnight culture in YPD media supplemented with 4 mM oleic acid, cells of strain H99 exhibited higher mean and maximum fluorescence intensity of neutral lipid. **F)** After overnight culture in YPD media supplemented with 4 mM oleic acid, cells of strain 409 exhibited no significant change in mean fluorescence intensity of neutral lipid but had a higher maximum fluorescence intensity.

### Proposed model of interaction of cryptococci with natural aqueous environments

We propose that cryptococcal cells in terrestrial reservoirs can be carried by freshwater into marine environments, where layering of freshwater over seawater results in a halocline interface that keeps cryptococci suspended at the water surface (**Figure 8**). Polysaccharide rafts would further prolong cell settling and enhance adherence to debris or biofilm formation. In the absence of a halocline, the rate of cell settling is a function of the cell’s gravity and the salinity of the water, with higher salinity water contributing more buoyant force.

**Figure 8.**
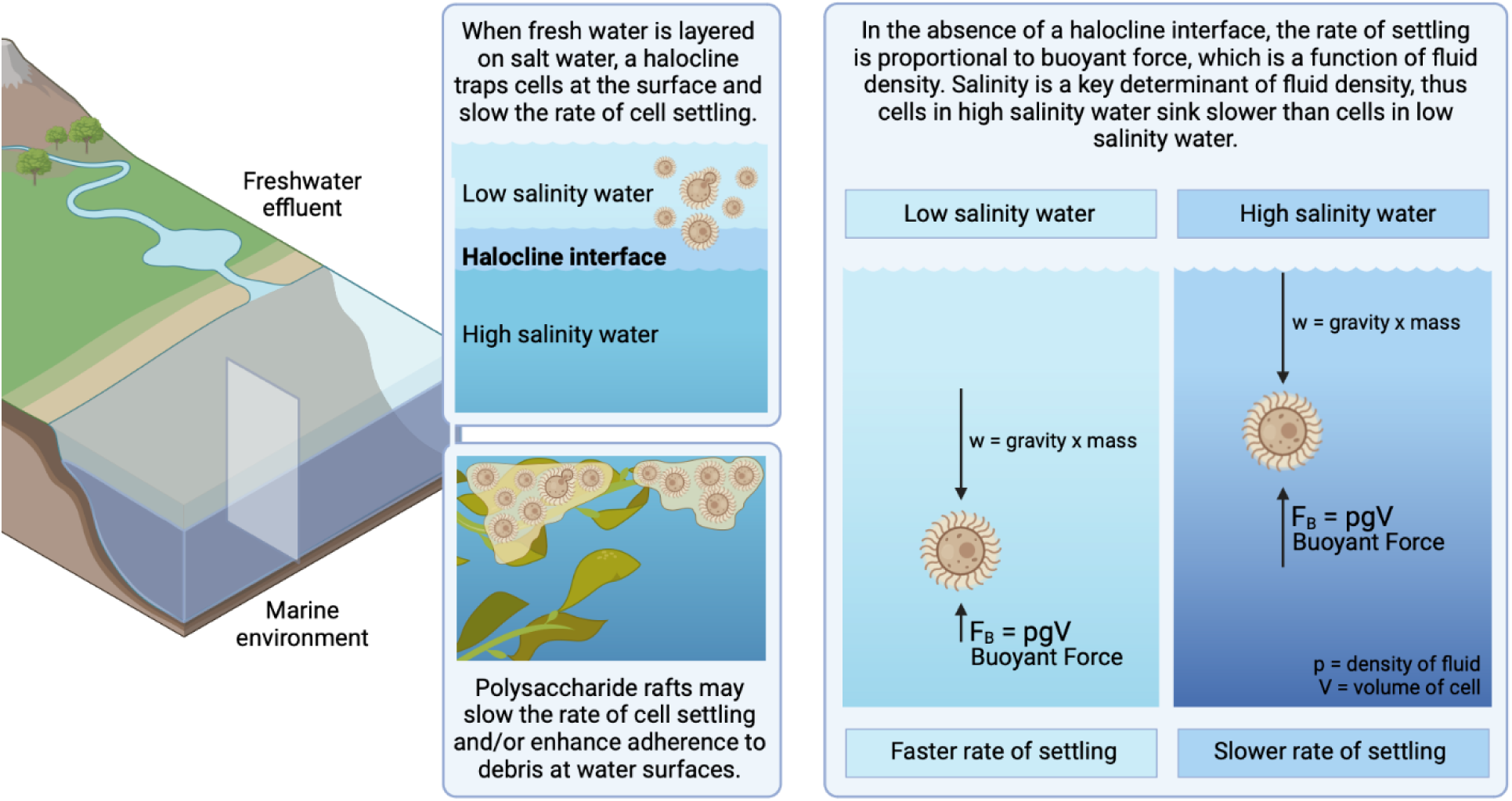
Proposed model of interaction of cryptococci in natural aqueous environments. We propose that cryptococcal cells residing in terrestrial reservoirs, such as trees, soil, and avian guano, are carried by freshwater effluents into marine environments. The layering of freshwater over seawater results in a halocline interface, suspending cryptococci near the water surface and slowing the rate of cell settling. In the absence of a halocline, the rate of cell settling is a function of mass, gravity and the salinity of the water, with higher salinity water contributing more buoyant force. Thus, marine environments slow the rate of cell settling and may enable long-range cryptococcal transport on ocean currents. Polysaccharide rafts, being less dense than cryptococcal cells, may also slow the rate settling of entrapped cells and/or enhance adherence to debris at water surfaces. Created with BioRender.com.

## Discussion

The ecological niche for pathogenic cryptococci is thought to be primarily land-based, with both *C. neoformans* and *C. gattii* found in soil and tree hollows, and *C. neoformans* additionally found in association with avian guano. Inhalation of dry aerosolized spores or desiccated yeasts from these terrestrial reservoirs is the primary documented mode of infection. However, terrestrial reservoirs for cryptococci are also exposed to rain, agricultural runoff, and wind, and cells could thus be carried to aquatic environments. Wildfire smoke, for instance, has been shown to transport viable microbes, including fungi.^45–48^ Kidd et al. (2007)^13^ showed that experimentally, *C. gattii* could survive for weeks in seawater and deionized water. The documented infections of marine mammals with *C. gattii* raise the potential of respiratory exposure to cryptococci through inhalation of cells suspended in liquid droplets or wet aerosol. Because marine mammals are intermittent breathers that hold inspired air in their lungs while underwater, their breathing pattern begins with rapid, forceful exhalation of spent air shortly after breaching the surface.^49–51^ Dolphins, for instance, expel up to 130 L/s of air^51^ at speeds of over 20 m/s, aerosolizing surface water in the process,^52^ before rapidly inhaling a mixture of air and spray. This presents an opportunity for respiratory exposure to pathogens carried within the water column. Given that water may also play a role in maintaining ecological cycles involved in cryptococcal survival and dissemination, it is important to study mechanisms that contribute to cryptococcal persistence in aqueous environments.

Few studies have evaluated cellular structures and variables affecting aqueous transport of cryptococci. Vij et al. (2018) demonstrated that cryptococci with large capsules had lower cellular density and cells without capsules had higher density, suggesting that the capsule could confer buoyancy and facilitate aqueous transport.^33^ Multiple findings in the present study confirm this role for the capsule. The rate of passive settling varied significantly by strain and correlated with cell densities, with cells of the *cap59* acapsular mutant sinking most rapidly, followed by strains H99, WM161, and finally 409. Strain-specific differences in density corresponded to different cell measurements, with *cap59* having a larger cell body and no capsule, while strain 409 had a larger capsule:body volume ratio. We also observed a higher baseline lipid content in strain 409 compared to H99, which could further contribute to strain 409’s lower cell density.

Capsular polysaccharides are highly hydrophilic and the capsule is highly intercalated with water, forming a hydrated shell around the cell body.^53^ The *C. neoformans* capsule has negatively charged glucuronic acid groups that bind divalent cations^54^ and contribute to repulsion of cells.^55^ Conversely, acapsular cells are notoriously clumpy when examined microscopically,^55^ a property that would accelerate settling^56^ and which we observed was enhanced in the presence of seawater, with macroscopic clumps of *cap59* cells adhering to the cuvette. The outer surface of the capsule is also hydrophobic,^57^ which may keep encapsulated cryptococci spaced apart as they settle through water, further contributing to cell suspension. On immunocytochemistry, capsules of *C. gattii* strains WM161 and 409 were also more diffuse and less compacted than capsules of H99 grown under the same conditions; differences in the structure of the capsule could also affect cell settling.

Cryptococcal capsule growth was not significantly induced by short-term incubation in seawater, supporting our hypothesis that capsule induction is not the sole mechanism by which cryptococci modulate buoyancy in natural environments. However, both *C. gattii* strains manifested significantly larger capsules after incubation in PBS; this was not observed in *C. neoformans* strain H99. Capsular growth is a response to cellular stress, such as in nutrient-poor environments. If *C. gattii* strains more readily form capsule in response to low salinity water, this could be an additional strategy to maintain buoyancy in freshwater environments.

We observed an additional phenomenon contributing to strain-specific cryptococcal cell buoyancy: polysaccharide raft formation. *C. neoformans* and *C. gattii* secrete exopolysaccharide (EPS) during culture and infection.^58, 59^ In this study, at various times of passive settling, all encapsulated stains developed a translucent upper region; this was not observed for the *cap59* acapsular mutant, consistent with prior work suggesting the *CAP59* gene is essential for polysaccharide export.^36^ This finding was most pronounced for strain 409, in which a large distinct upper layer was visible after over 24 h. On microscopic evaluation, this layer contained copious acellular material interspersed with cells with a high capsule:body volume ratio, while the lower layer was densely packed with cells at approximately 100x higher concentration. Our results support that this material is largely comprised of glucuronoxylomannan (GXM) polysaccharide, which is also the principal component of the cryptococcal capsule. Diluting overnight cultures with fresh YPD media accelerated settling of all encapsulated strains but did not affect the settling of *cap59*, demonstrating that EPS influences buoyancy in a dose-dependent manner. We hypothesize that polysaccharide secreted or shed during growth of encapsulated strains could contribute to buoyancy by remaining near the water surface and acting as a raft for entrapped cells. EPS from *C. laurentii* was reported to facilitate and stabilize oil-water emulsions and to increase viscosity and drag, both of which would slow the rate of cell settling.^60^ The discovery of polysaccharide rafts that aid in flotation suggest a new role for EPS in promoting aqueous transport.

Different laboratory methods of EPS isolation have varying effects on polysaccharide organization, structure, and aggregation.^58, 61, 62^ In this study, passive settling of a culture and collection of the translucent upper layer enriched the sample for EPS compared to direct sampling of a confluent culture. This method may supplement existing EPS collection techniques. Strain-specific differences in settling time should be considered when determining the optimal time to sample this layer, and may reflect differences in amount, composition, or structure of EPS. High concentrations of EPS also appeared to inhibit binding of 18B7 mAb to cells trapped within EPS aggregates. The role of EPS in sequestering cells from antibody may be an immune evasion mechanism, given the importance of antibody-mediated opsonization in the response to cryptococcal infection. Aggregates of polysaccharide and cells have been described during *in vitro* infection of macrophages with *C. neoformans* or *C. gattii*^59^ and in the context of cryptococcal biofilm formation.^63–65^ Our methods also preserved macromolecular structures, allowing visualization of relationships between polysaccharide aggregates and entrapped cells, which may be applicable to future studies of biofilm formation.

To test the hypothesis that freshwater could carry cryptococci from land to the air-water surface, we experimentally replicated the ecological phenomenon of halocline formation, in which low salinity water forms a stable layer above seawater. In nature, particles traverse the halocline as a function of their density and can become suspended at this interface, creating a unique composition of nutrients, debris, and microbes.^66^ In the presence of a halocline, encapsulated cryptococci were trapped in the upper 1 cm of the water column for over 60 min. When no halocline was present, cells grown under identical culture conditions rapidly sank past the air-water interface. The effect of the halocline on cell suspension was consistent across all four strains tested, including the *cap59* mutant, implying that this effect is independent of the capsule. Cells remained suspended at the air-water interface even in the presence of movement, suggesting that cryptococci could remain suspended while carried by natural water currents. Higher density fluids confer more buoyant force; in the absence of a halocline, cells indeed settled slower in higher salinity media than in lower salinity media.

In this study, we observed strain-specific differences in cell density, capsule:body volume ratio, and polysaccharide production that affected cell settling, and demonstrated that halocline formation enhances buoyancy. By increasing persistence in surface water, cryptococci are more likely to be carried by waves to new environmental niches, encounter debris upon which to form biofilms, and encounter susceptible hosts. Our results identify *Cryptococcus* spp. characteristics that affect buoyancy and support the view that this fungus can survive, persist, and be transported in aqueous environments.

## Acknowledgements

Microscopy images were completed using the Light Microscopy Core of the Department of Molecular Microbiology and Immunology at the Johns Hopkins Bloomberg School of Public Health. We thank Shana Lee for performing the mucicarmine cytologic stain, and Jeff Cimprich for the three-dimensional render of the capsule in Figure 1B. Support for I.A.J. was provided by NIH T32 OD011089. A.C. was supported in part by NIH R01grants AI052733-16, AI152078-01, and HL059842-19.

## Supplemental Tables

**Supplemental Table 1:** Concentrations and specific gravity of sodium chloride solutions.

| [NaCl] (g/L) | Molarity (M) | Specific gravity |
| --- | --- | --- |
| 1.5 | 0.026 | 1.002 |
| 3 | 0.051 | 1.003 |
| 9.5 | 0.163 | 1.007 |
| 21.2 | 0.363 | 1.016 |
| 39.5 | 0.676 | 1.028 |
| 88 | 1.506 | 1.060 |

---

**Supplemental Figure 1.**
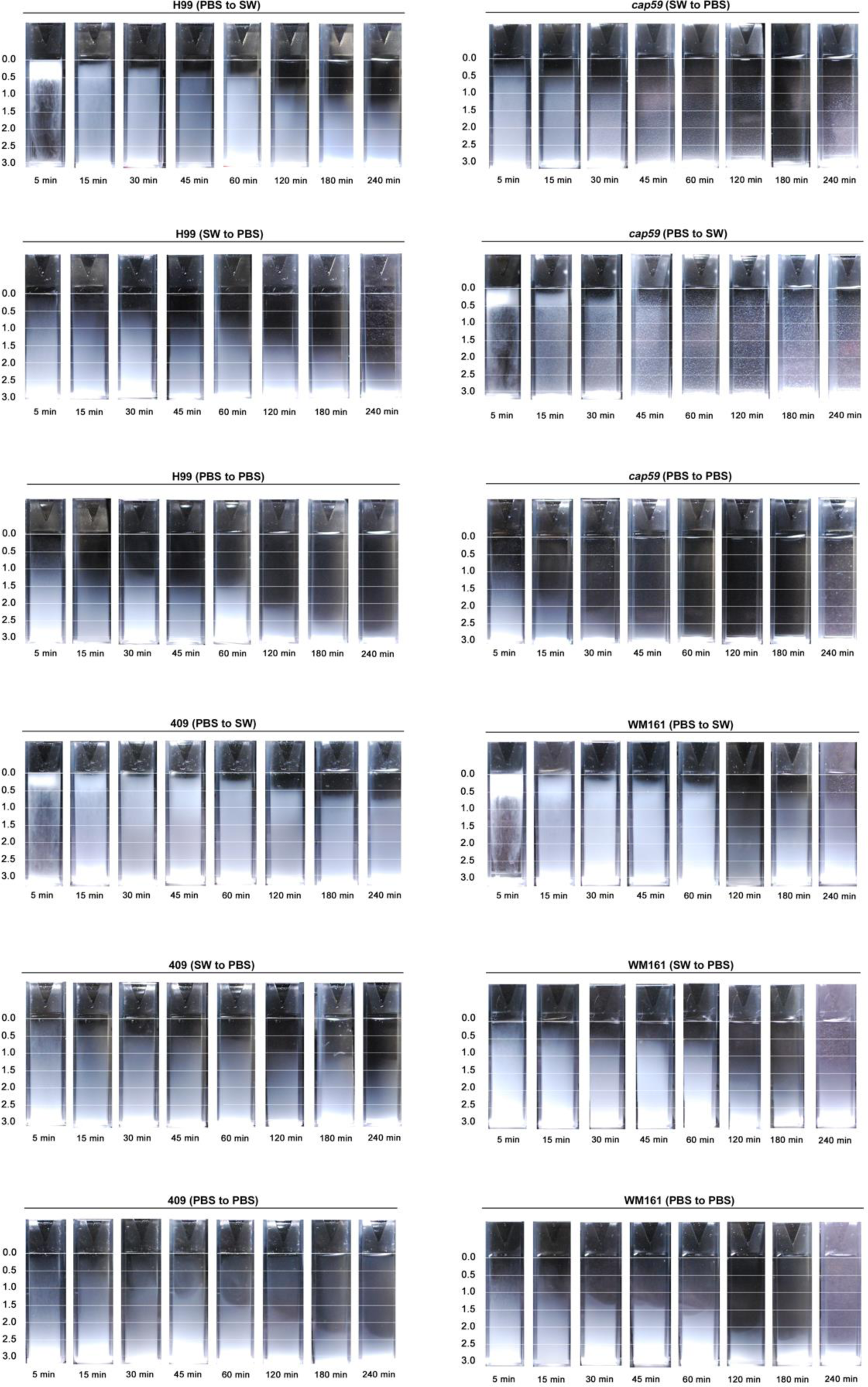

**Supplemental Figure 2.**
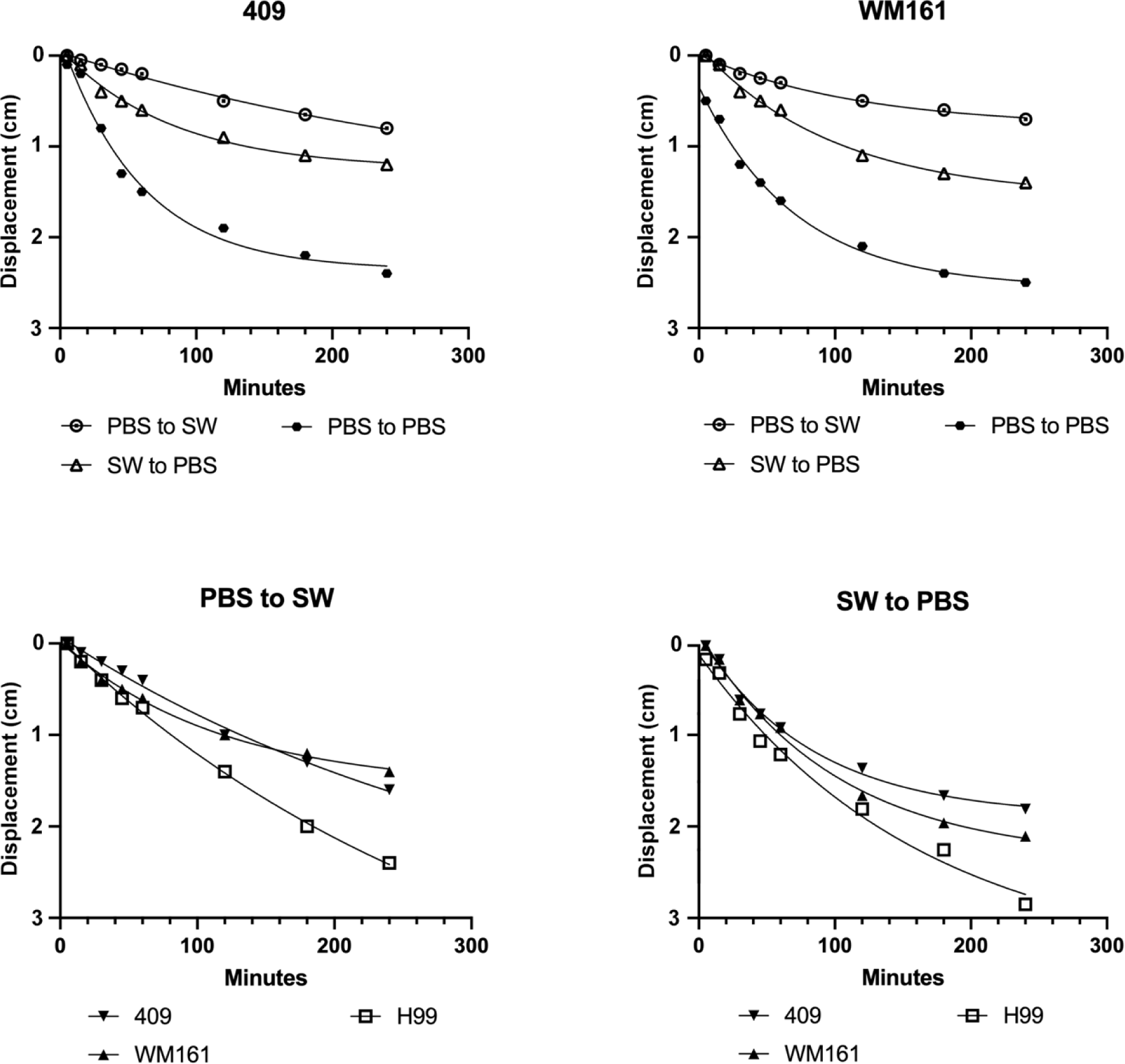

**Supplemental Figure 3.**
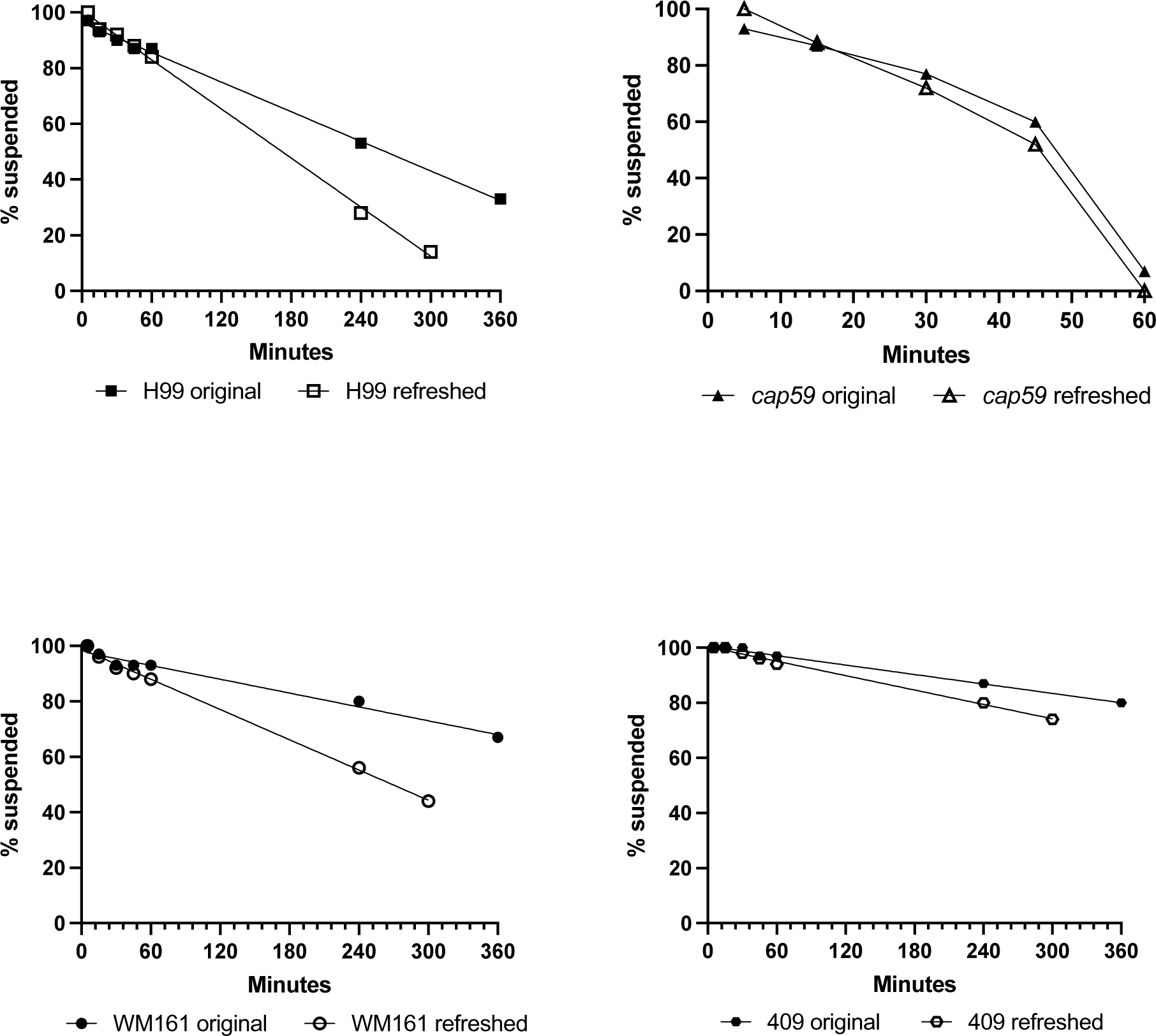

